# Multiplexed triage of candidate biomarkers in plasma using internal standard triggered-parallel reaction monitoring mass spectrometry

**DOI:** 10.1101/2021.09.02.458791

**Authors:** Jacob J. Kennedy, Jeffrey R. Whiteaker, Richard G. Ivey, Aura Burian, Shrabanti Chowdhury, Chia-Feng Tsai, Tao Liu, ChenWei Lin, Oscar Murillo, Rachel Lundeen, Lisa A. Jones, Philip R. Gafken, Gary Longton, Karin D. Rodland, Steven Skates, John Landua, Pei Wang, Michael T. Lewis, Amanda G. Paulovich

## Abstract

Despite advances in proteomic technologies, clinical translation of plasma biomarkers remains low, partly due to a major bottleneck between the discovery of candidate biomarkers and downstream costly clinical validation studies. Due to a dearth of multiplexable assays, generally only a few candidate biomarkers are tested, and the validation success rate is accordingly low. Here, we demonstrate the capability of internal standard triggered-parallel reaction monitoring (IS-PRM) to prioritize candidate biomarkers for validation studies. A 5,176-plex assay coupling immunodepletion and fractionation with IS-PRM was developed and implemented in human plasma to quantify peptides representing 1,314 breast cancer biomarker candidates. Compared to prior approaches using data-dependent analysis, IS-PRM showed improved sensitivity (912 vs 295 proteins quantified) and precision (CV 0.1 vs 0.27) enabling rank-ordering of candidate biomarkers for validation studies. The assay greatly expands capabilities for quantification of large numbers of proteins and is well suited for prioritization of viable candidate biomarkers.

## Introduction

Blood plasma is an easily accessed biofluid that reflects the physiological state of a patient; thus, it remains an attractive source of clinical biomarkers^1,2^. Despite considerable investment and advances in liquid chromatography mass spectrometry-based (LC-MS/MS) proteomic technologies that allow for deep coverage and quantification of proteins^3,4^, translation of biomarker discoveries to clinical use remains slow, tedious, and generally disappointing^5,6^. A large factor contributing to this state of the field is the mismatch between the large number of potential biomarkers identified and the resources required for their validation. A method to prioritize amongst candidate biomarkers to identify those with the greatest probability of clinical utility would allow clinical validation efforts to focus on the subset of candidates most likely to succeed^7^.

The emergence of targeted mass spectrometry-based proteomics approaches (e.g., multiple reaction monitoring (MRM) and parallel reaction monitoring (PRM)^8–10^) enables highly sensitive, specific, and multiplexable assays that can be implemented with relatively low cost (compared to traditional immunoassays)^11^. These approaches have been incorporated into pipelines for biomarker evaluation^7,12–14^ and have been applied to biomarker development efforts^15,16^. However, even with optimized parameters and careful attention to method details (e.g., tight retention time windows, elimination of overlapping interfering transitions, enrichment and/or fractionation for low abundance targets), it is a challenge to measure more than a few hundred proteins and maintain high analytical performance using these approaches^17–20^. Thus, there is a need for a method capable of measuring a thousand or more biomarker candidates in plasma with high sensitivity and specificity. Such an approach could be used in the biomarker development pipeline to triage a large list of candidates down to a smaller number that can be applied with more quantitative rigor at increased throughput.

The recent development of internal standard triggered-parallel reaction monitoring mass spectrometry^21^ (IS-PRM-MS, or SureQuant) has allowed for high multiplexing with the benefits of the performance of PRM^22,23^. The IS-PRM method expands the capacity of the PRM method without relying on retention time windows or co-isolation of target peptides by instead relying on added internal standards to trigger the real-time measurement of endogenous peptides. Upon detection of the internal standard, quantification is performed by PRM, allowing for highly sensitive and specific measurements.

In this study, we evaluated the effectiveness of IS-PRM in the context of prioritizing candidate biomarkers for more costly validation studies. As a test case, we developed an IS-PRM method to quantify 5,176 peptides representing 1,314 candidate breast cancer plasma biomarker proteins identified using a novel strategy leveraging preclinical patient derived xenograft (PDX) mouse models. We hypothesized that the IS-PRM method could quantify these candidates in human plasma with high specificity and precision to enable the rank ordering of candidate biomarkers for further investment of resources to perform validation studies in large patient cohorts. The methodology developed herein presents a significant advance in reliable quantification and verification of large numbers of plasma-based biomarker candidates, and the approach is generally applicable to other diseases or translational studies requiring highly precise relative quantification of large sets of proteins.

## Results

### Overview

We evaluated the utility of a highly multiplexed IS-PRM assay for prioritizing a large list of candidate plasma biomarkers of breast cancer (identified in breast cancer PDX models; see **Supplementary Fig. 1**) for resource-intensive follow up clinical validation studies. The IS-PRM (**Fig. 1**) assay quantifies endogenous (“light”) peptide after first observing and identifying its cognate spiked-in isotope-labeled (“heavy”) internal standard peptide. After a positive identification is confirmed, quantification is achieved by performing targeted PRM on the endogenous peptide. This method accomplishes high sensitivity and specificity with improved multiplexing (required to triage large numbers of biomarker candidates) by using fast MS scans for identification and maximizing the time devoted to quantitative scans. In addition, the inclusion lists employed by the method can survey for tens of thousands of target precursors making the method easier to implement because it does not require characterization and monitoring of retention time windows.

**Fig. 1:**
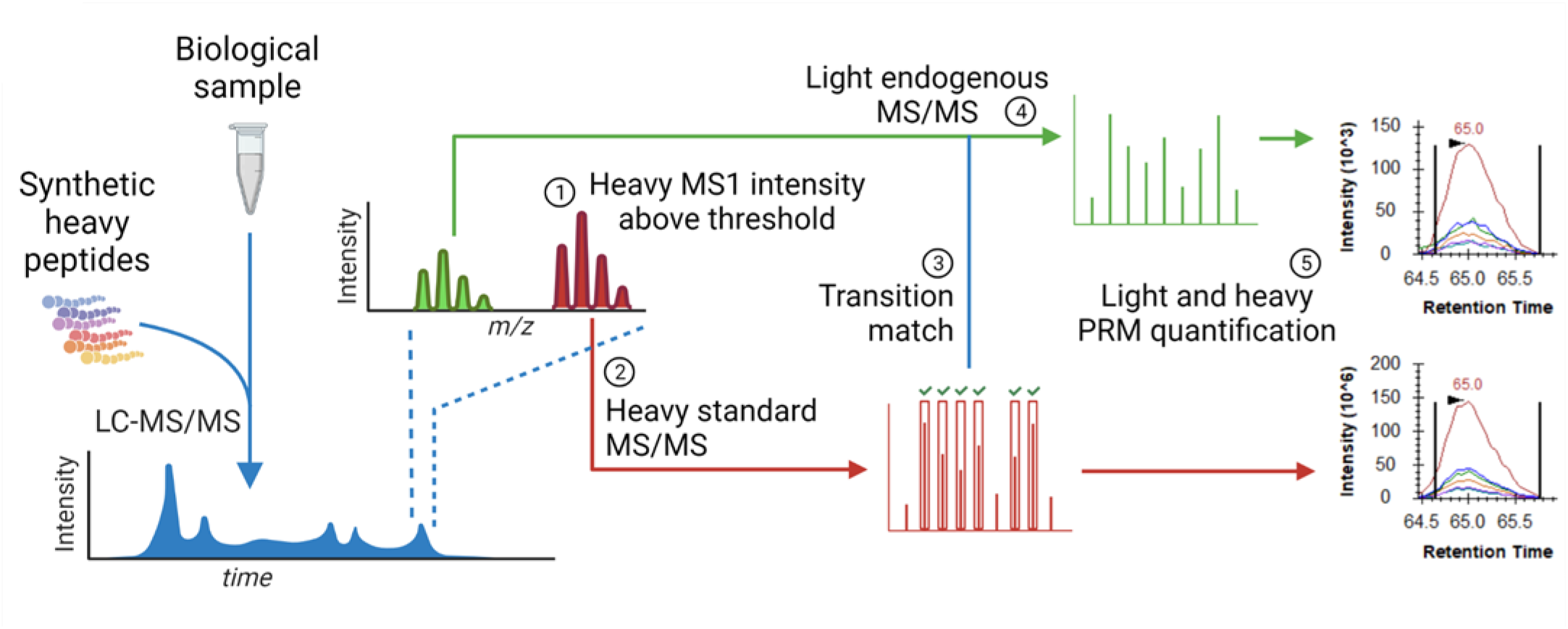
A targeted MS workflow based on IS-PRM applied to prioritization of breast cancer biomarkers for validation studies. **a** Proteins in a biological sample are converted to peptides by enzymatic digestion and synthetic heavy isotope-labeled standards for each target peptide are spiked into the mixture. In step 1, the peptide mixture is analyzed by liquid chromatography-mass spectrometry, using an inclusion list containing one precursor (peptide) and six transition (fragment) m/z associated with each heavy standard. An MS1 scan is performed to look for precursor m/z on the inclusion list. Once a heavy precursor m/z has been observed above a given intensity threshold, step 2 is initiated, where a low-resolution MS2 scan is triggered to identify transition fragment ions associated with the targeted heavy peptide. If 5 out of 6 transitions are observed, then a high-resolution MS2 scan is initiated for the light version of the peptide in step 3. Finally, in step 4, parallel reaction monitoring (PRM) using the high resolution MS2 scans of light and heavy peptides allows for conclusive identification of the peptide sequence and relative quantification of the endogenous “light” peptide, which is reported as the peak area ratio (PAR) of the light peptide intensity over the heavy peptide intensity. **b** Candidate protein biomarkers were identified by LC-MS/MS analyses of plasma from mice harboring human patient-derived xenografts (PDX) of breast cancer to identify *human* proteins secreted, leaked, or shed from tumors (5,498 unique human-specific peptides, mapping to 1,314 human proteins). As a test case for prioritizing the candidates for further up validation studies, the IS-PRM assay was used to interrogate differential expression in plasma from breast cancer patients versus benign controls. 138 human plasma samples were individually depleted of high- and mid-abundance proteins and combined accordingly to create cancer and control pools. The heavy isotope-labeled standard peptide mix was spiked into the digested pools prior to fractionation by basic reverse phase (bRP) liquid chromatography. Each of the 24 bRP fractions was subjected to the IS-PRM method using a Thermo Orbitrap Eclipse Tribrid mass spectrometer. The peak area ratio of endogenous light peptides relative to heavy spiked peptide was used to quantify the relative expression between pools.

As a test case for employing IS-PRM in triaging biomarker candidates, we used the method to rank order a list of 1,314 candidate biomarker proteins (**Supplementary Fig. 1**) by quantifying the differences in expression in pooled plasma samples from women diagnosed with breast cancer vs plasma samples from women diagnosed with benign breast lesions. The 1,314 breast cancer biomarker candidates were identified as *human* proteins in the plasma of patient-derived xenograft (PDX)-bearing mouse breast cancer models, where proteins leaked, secreted, or shed from the transplanted human breast tumors were the exclusive source of human proteins in the plasma. Of note, 1,179 (90%) of the candidate biomarkers were previously observed in proteomic profiles of human breast cancers^24^.

### IS-PRM method development and characterization

Up to 3 proteotypic peptides representing each of the 1,314 candidate biomarker proteins were identified from amongst peptides empirically observed in the PDX plasma biomarker discovery data (n=2852), supplemented by additional peptides from Peptide Atlas^25^ (http://www.peptideatlas.org/) and/or SRMAtlas^26^ (http://www.srmatlas.org/) (n=2122) and peptides identified from previous assay development efforts (n=208)^27^. After filtering peptides based on length (between 7 and 25 amino acids) and hydrophobicity (SSRCalc^28^ between 10-40), we identified a total of 5,176 peptides, with ≥3 peptides identified for 1,303 (99%) candidate biomarker proteins (**Supplementary Fig. 2**).

Heavy stable isotope-labeled standard peptides (SIS) were synthesized for each target peptide (**Supplementary Table 1**) and used to determine the optimum precursor m/z for the IS-PRM inclusion list, the fragment ions to be used for peptide identification, and the intensity thresholds for triggering the identification scan. To optimize these parameters, a mixture of the 5,176 heavy peptides (~500 fmol) was spiked into 200 μg of trypsin/LysC-proteolyzed yeast lysate. The mixture was fractionated by basic reverse phase (bRP) liquid chromatography into 12 fractions. Each fraction was then subjected to data dependent acquisition (DDA) LC-MS/MS using an inclusion list (i.e., directed DDA) containing m/z values for +2 and +3 charge states for each SIS peptide. Spectra matching the heavy peptide sequences were used to identify the six most intense fragment ions for each precursor, and to identify the precursor charge state that had the most intense sum of these six transitions. Using these empirical data, we set the intensity threshold for triggering the identification scan at 2% of the maximum height of the MS1 intensity of the precursor to maintain the highest sensitivity without triggering on noise. The precursor m/z and intensity thresholds are listed in **Supplementary Table 2**, and fragment ions used for identification and quantification are listed in **Supplementary Table 3**.

The analytical performance of the IS-PRM method was characterized using a response curve consisting of a ten-fold serial dilution of human cell (MCF10A) lysate into yeast lysate (100% MCF10A to 0.1%, blanks were prepared using 0% MCF10A). The MCF10A concentration levels corresponded to an approximate MCF10A cell count of 200,000 to 200 cells, respectively. Each concentration point underwent proteolytic digestion, addition of SIS peptides, and separation into 6 bRP fractions. Each fraction was analyzed by the IS-PRM method using triplicate injections on the LC-MS (**Fig. 2a**). For a peptide to be classified as quantified, we imposed the following requirements which had to be satisfied in at least two of the three replicates: (i) at least 4 transitions (light endogenous peptides) or 5 transitions (heavy SIS peptides) were present in the MS2 spectra, (ii) the ratio dot product of MS2 spectra from heavy and light peptides was > 0.98, (iii) at least 5 points across the peak were profiled in the chromatogram, and (iv) the peak area was > 5,000. All integrations were manually checked, and 93 (~2%) peptides had a fragment ion with interference in either the heavy or light peptide removed from the analysis.

**Fig. 2:**
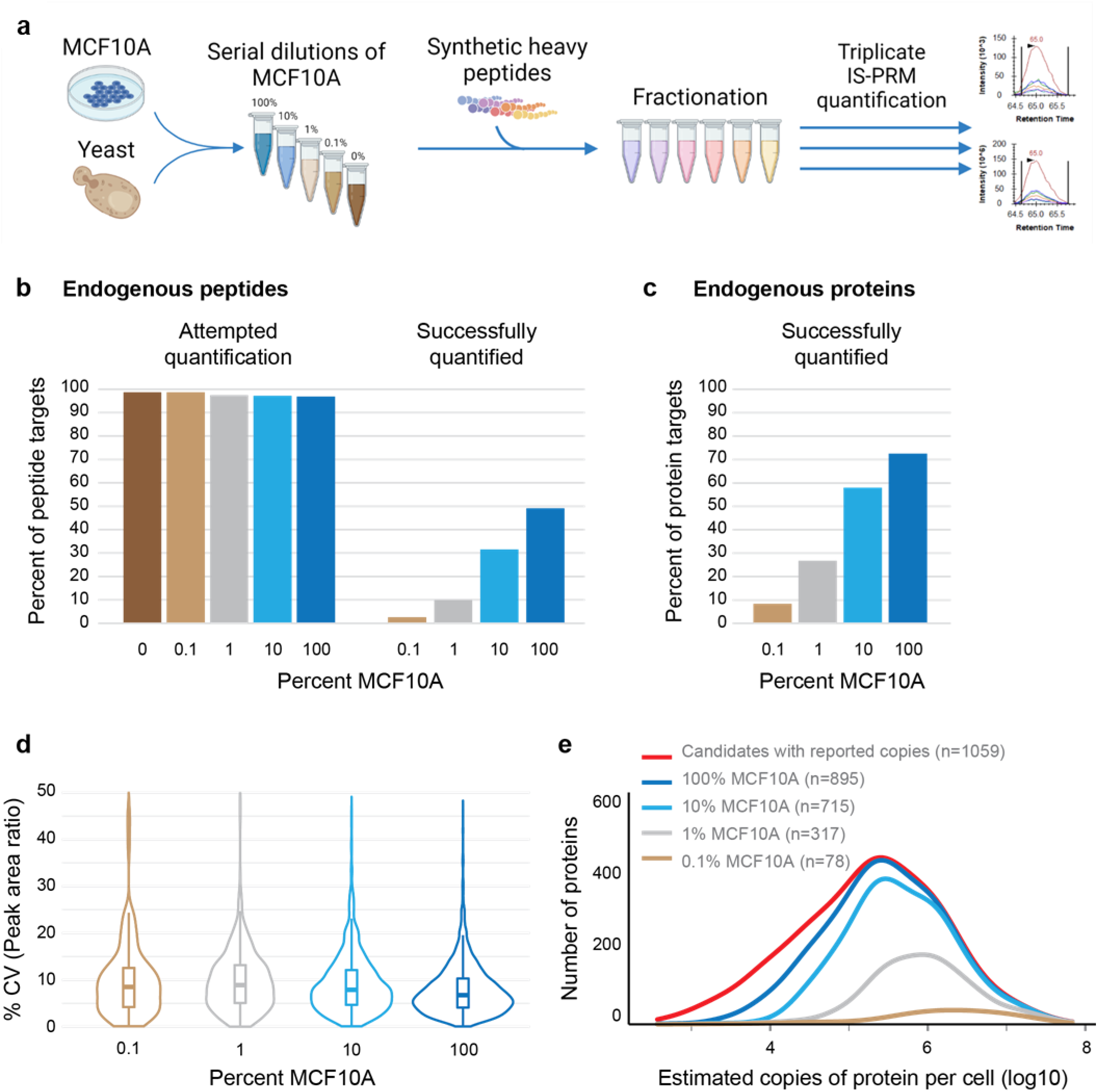
Characterization of the analytical performance of the IS-PRM assay. **a** Digested MCF10A cell lysate was serially diluted in digested yeast lysate to concentration points of 100%, 10%, 1% and 0.1% MCF10A. A 0% blank was also included. 50 μg of each point was proteolytically digested, spiked with heavy standard peptides to 1,314 proteins, and fractionated into six samples, each of which were analyzed in triplicate by the optimized IS-PRM method. **b** The percent of peptides triggered for quantification and successfully quantified is plotted for each concentration sample. Light peptides triggered for quantification refers to those peptides meeting the detection threshold and fragment ion requirement for measurement. Endogenous peptides successfully quantified refers to those peptides meeting all quantification criteria. **c** Percent of total protein targets quantified. **d** Violin plot showing the precision of the replicate measurements of the heavy to light peak area ratios for each dilution point. Bold line shows median, box shows inner quartile, vertical line shows 5-95 percentile, outliers are shown as individual points, density of measurements is indicated by the thin line. **e** Histogram showing the distribution of number of proteins detected according to the protein level per cell. The majority of the target proteins were quantified in the response curve. The number of proteins detected for each dilution point is indicated in the legend.

Response curve results for the IS-PRM assay are summarized in **Fig. 2** and data are provided in **Supplementary Table 4**. The IS-PRM method triggered the quantification of most of the light peptides (**Fig. 2b**). Endogenous signals were quantified for nearly half of the peptides (n=2,443) at the highest concentration of MCF10A (**Fig. 2b**), resulting in quantification of 953 proteins (73%) of the targeted proteins (**Fig. 2c**). Decreasing the percentage of MCF10A cells resulted in the expected decrease in proteins quantified. The method exhibited excellent analytical precision, with a median coefficient of variation (CV) of 7.7% across all concentration points (**Fig. 2d**). To estimate the sensitivity of the method for detection of low abundance proteins, we used the number of proteins expressed per cell reported in Ly et. al.^29^. **Fig. 2e** shows a histogram of proteins detected by the IS-PRM method in each dilution point versus the number of proteins per cell. As expected, as the MCF10A cells were diluted, the histogram curve shifts to those proteins that were most abundant.

We next compared the sensitivity and precision of the IS-PRM method to directed DDA, which is also capable of highly multiplexed identification and quantification and which has been previously used to prioritize candidate biomarkers^7,30^. The target peptides were measured using both IS-PRM and directed DDA in a common trypsin/LysC-proteolyzed pool of human plasma depleted of abundant proteins and fractionated by bRP chromatography in the same manner as the yeast sample above (**Fig. 3a**). While both directed DDA and IS-PRM use an inclusion list to target precursors for MS2 fragmentation, the method of quantification differs considerably. Directed DDA uses the MS1 scans to create an extracted ion chromatogram for quantification, whereas IS-PRM quantifies the target peptide based on the intensities of selected fragment ions in the MS2 scan. The IS-PRM assay triggered quantification of more light peptides (4674 vs 3991) and exhibited better sensitivity (**Fig. 3b)** compared to directed DDA, as shown by quantifying 1683 endogenous peptides in IS-PRM versus 436 in directed DDA. The difference in peptide quantification rate was even more pronounced at the protein level (**Fig. 3c**), where IS-PRM quantified 912 proteins versus 295 for directed DDA. The precision of the approaches was estimated by calculating the CV from peak area ratios measured in the neighboring bRP fraction as technical replicates of the LC-MS/MS measurements. As expected, the median CV was lower in results from the IS-PRM analysis compared to the directed DDA analysis (0.10 versus 0.27; **Fig. 3d**). This is a result of improved signal-to-noise and reduction in interferences in using the MS2 signals for quantification in PRM compared to MS1 signals in directed DDA.

**Fig. 3:**
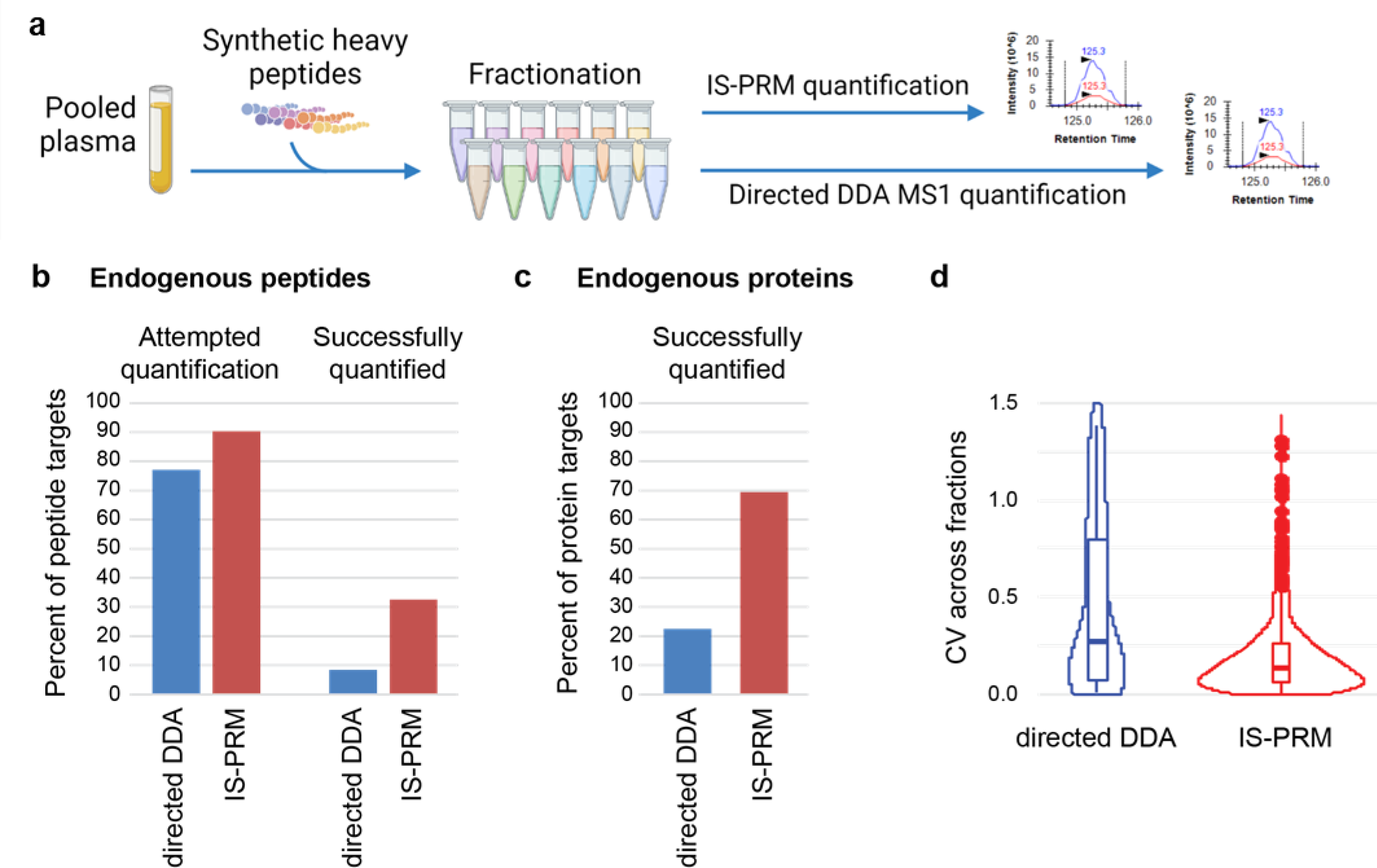
The IS-PRM assay improves relative quantification of proteins compared to directed DDA methods. **a** Comparison of the IS-PRM assay to directed DDA was conducted in a pool of depleted plasma that was proteolytically digested, spiked with 5,176 heavy standard peptides to 1,314 proteins, and separated into 12 bRP fractions, each of which were analyzed by both the IS-PRM assay as well as by directed DDA. **b** The percent of total targeted peptides is plotted for targeted and quantified endogenous peptides for each acquisition method. ‘Attempted quantification’ refers to light peptides triggered for quantification after meeting the detection threshold and fragment ion requirement for measurement. A total of 4674 (IS-PRM) and 3991 (directed DDA) peptides were triggered. ‘Successfully quantified’ refers to endogenous peptides successfully quantified and meeting all quantification criteria. A total of 1683 (IS-PRM) and 436 (directed DDA) peptides were successfully quantified. **c** Percent of total protein targets quantified, total number of proteins quantified were 912 (IS-PRM) and 295 (directed DDA). **d** Violin plot showing distribution of standard error for endogenous peptide measurements measured by using the PAR in neighboring bRP fractions as technical replicates. Bold line shows median, box shows inner quartile, vertical line shows 5-95 percentile, outliers are shown as individual points, density of measurements is indicated by the thin line.

### Evaluation of the IS-PRM method for highly multiplexed quantification of candidate biomarkers in human plasma

We next applied the 5,176-plex IS-PRM assay to quantify the biomarker candidates in 3 pools of human plasmas from women diagnosed with breast cancer and 3 pools of human plasmas from women diagnosed with benign breast lesions (**Fig. 4a**), with a goal of rank-ordering the 1,314 candidate biomarkers to identify those meriting further evaluation in larger, case-control validation studies. Each pool represented 19-20 women. Two hundred micrograms of each plasma pool underwent reduction, alkylation, and proteolytic digestion. The digested plasma pools were desalted, spiked with all 5,176 SIS peptides (~500 fmol) and fractionated into 24 fractions using bRP chromatography.

**Fig. 4:**
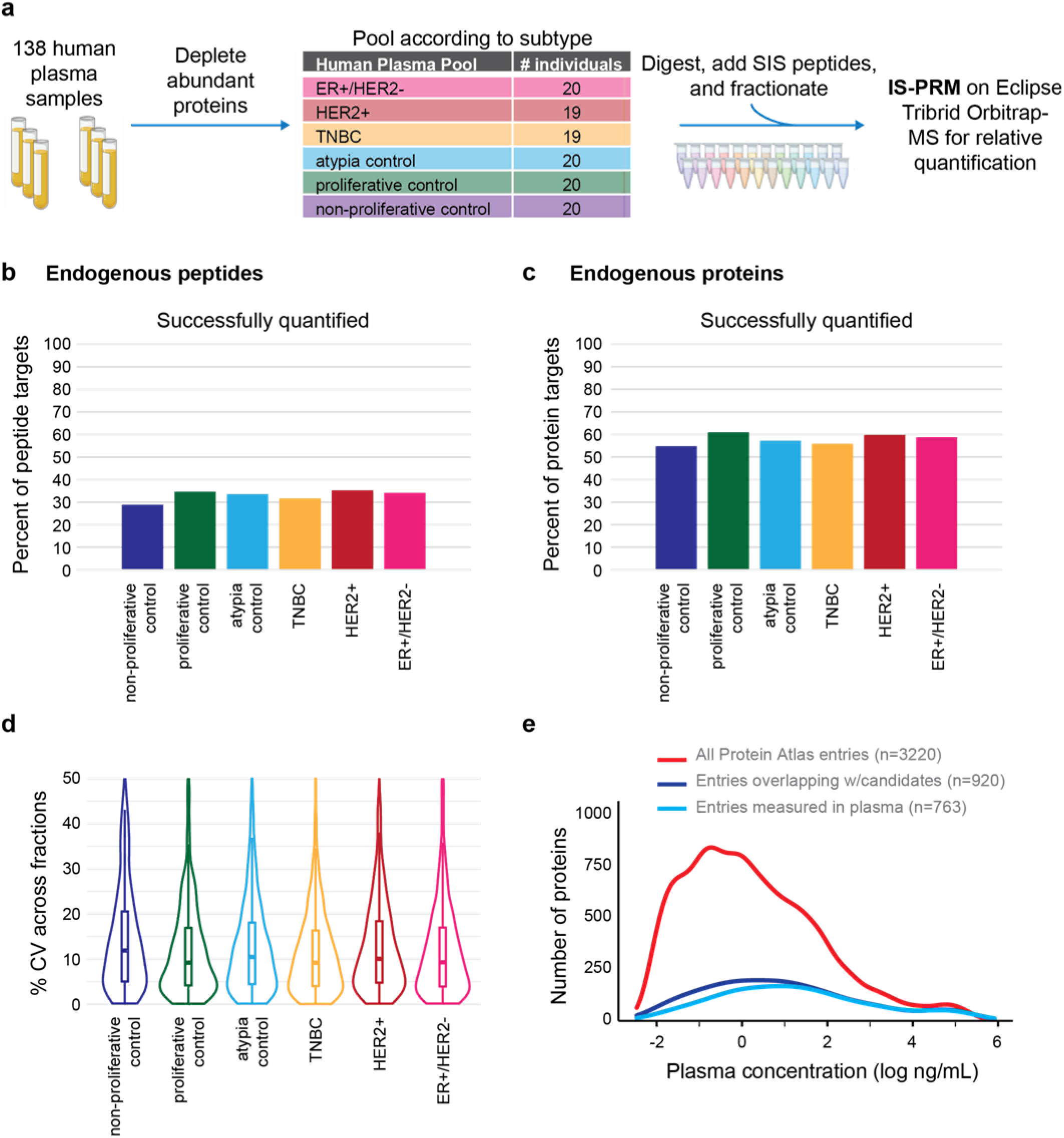
Applying the IS-PRM assay to prioritize the biomarker candidate proteins in plasma of human breast cancer patients. **a** 138 plasmas were depleted of high- and mid-abundance proteins and pooled into 6 samples (3 breast cancer subtype pools and 3 benign breast lesion controls), proteolytically digested, spiked with heavy standard peptides to 1,314 proteins, and fractionated into 24 samples, each of which were analyzed by the optimized IS-PRM method. **b** The percent of endogenous light signals meeting quantification criteria in each of the plasma pools. **c** The percent of candidate protein biomarkers with endogenous levels measured in each of the plasma pools. **d** Violin box plot showing distribution of standard error for endogenous peptide measurements measured by using the PAR in neighboring bRP fractions as technical replicates. Bold line shows median, box shows inner quartile, vertical line shows 5-95 percentile, outliers are shown as individual points, density of measurements is indicated by the thin line. The precision of the measurements was measured as the standard error of the endogenous signal quantified in adjacent fractions. **e** Distribution of the number of proteins detected according to reported plasma concentration. The majority of the target proteins were quantified in all plasma pools and the candidate biomarkers and endogenous measurements based on the IS-PRM method represent a wide concentration range in plasma.

The IS-PRM method was applied to each of the 24 fractions (per pool), using a rigorous QC program to avoid any system degradation during the analysis (**Supplementary Fig. 3**). All peak integrations were manually reviewed, and interferences removed from 193 (~4%) peptides. Summed transition areas and number of transitions and points per peak are reported in **Supplementary Table 5**. In addition to the quantification criteria used for the response curve (above), we required endogenous signal to be >2x the signal from blank runs (**Supplementary Table 6**). On average, endogenous signals were measured for 1,708 (33%) of the target peptides (**Fig. 4b**) across the pools, corresponding to 760 (58%) proteins (**Fig. 4c**). The sum of all proteins quantified across the plasma pools was 893 (68%). Analytical reproducibility was determined using the endogenous measurements in the neighboring bRP fraction as a technical replicate (n=2). **Fig. 4d** shows the distribution of CV for each plasma pool (median across all measurements = 11.0%). Using the plasma concentration for proteins reported in the Human Plasma Peptide Atlas^31^ and the median of multiple peptide measurements per protein, we estimated the range of plasma concentrations for the proteins quantified by the IS-PRM assay. **Fig 4e** shows the distribution of protein concentrations for the candidate biomarkers with concentrations extending to below the ng/mL level.

The overall distributions of the 893 candidate biomarker protein abundances across the six plasma pools, shown in **Fig. 5a**, varied widely. To determine if the candidate biomarker protein signals were higher in the cancer plasma pools, we tested the proteins for significant differences (p < 0.001) in each cancer pool compared to at least 2 of the confounding (i.e., benign) control plasma pools. To allow for variability in the control samples, we used a regression trend approach, which accounted for measurements that were lowest in the non-proliferative control, increasing in the proliferative and the atypia controls, and reaching a maximum in the cancer subtype sample (i.e., candidates whose plasma levels progressively increased as the biology of the breast lesions became more aggressive). An example of an individual protein featuring this trend is shown in **Fig. 5b,** the results for all proteins are reported in **Supplementary Table 7**. Two peptides for PZP show consistent relative quantification (**Fig. 5b**), where the lowest measurement is seen in the non-proliferative control, followed by the proliferative and atypia controls, and finally the TNBC cancer subtype sample shows the highest levels. Overall, there were 162 candidate proteins showing significant differences in at least one of the three cancer subtypes (triple negative, HER2 positive and ER positive/HER2 negative), and 22 were significant in all three (**Fig. 5c**). The distribution of measured abundances for the 22 overlapping proteins (**Fig. 5d**) reflects an improvement in differentiating the cancer pools from control (compared to total proteins measured). Compared to a random sampling of 22 proteins, the candidates overlapping from all three subtypes show a better differentiation from controls (**Fig. 5e**).

**Fig. 5:**
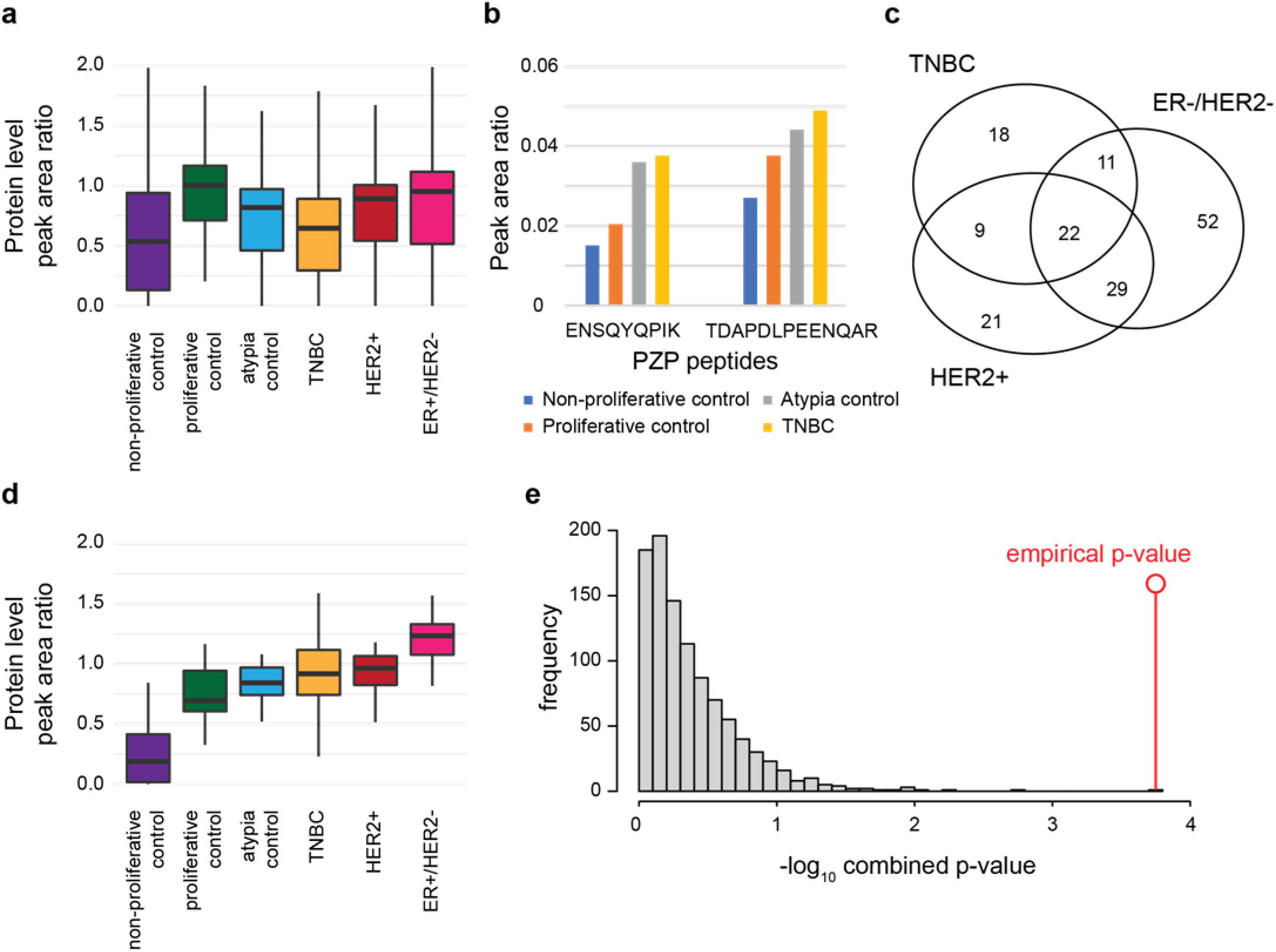
Verification of candidate biomarkers in the breast cancer plasma pools. **a** Box plot for proteins meeting quantification criteria in the depleted and fractionated plasma pools. The median of multiple peptide measurements was used for the protein value. Bold line shows median, box shows inner quartile, vertical line shows 5-95 percentile. **b** Plot of peak area ratio (light/heavy) for quantified peptides from PZP in three control pools and triple negative breast cancer (TNBC). An example of a candidate biomarker meeting significance testing in the TNBC breast cancer subtype with endogenous levels significantly higher (p < 0.001) in cancer compared to at least 2 of the 3 confounding control plasma pools with a significant regression trend (p < 0.01) (trend comparing non-proliferative control -> proliferative control -> atypia control -> cancer subtype). **c** Venn diagram of the 162 candidate biomarkers verified in the pooled case/control study. 22 of the candidates passed both cutoffs (p-value<0.001 and regression trend p-value <0.1) in all 3 breast cancer subtypes. **d** Box plot showing endogenous levels of the 22 proteins found higher in all three breast cancer subtypes compared to the three confounding control samples. Plotted as peak area ratio (PAR; light:heavy) using the median value from multiple measurements of peptides. Bold line shows median, box shows inner quartile, vertical line shows 5-95 percentile. **e** Histogram showing the combined p-value from differentiation between cancer subtypes and control plasmas using randomly sampled subsets of 22 proteins (1,000 permutations). The p-value for the set of overlapping 22 proteins verified by IS-PRM assay (p=0.00016) is shown by the red line. P-values are based on a student t-test.

## Discussion

With the application of modern approaches, many hundreds of candidate protein biomarkers of disease are readily identified, but few of these candidates are ever carried forward to clinical validation studies^32,33^, and almost none are clinically translated. This chasm between biomarker discovery and validation is largely caused by a paucity of validated multiplexable assays for quantification of protein biomarker candidates in clinical validation studies, leading to a largely arbitrary selection of a few candidate biomarkers for which commercial assays are available for advancement to clinical studies. An empirical method to rank order large lists of candidate biomarkers to identify those meritorious enough to warrant an investment in assay development and clinical validation could improve success rates for translating novel biomarkers into clinical use.

We demonstrate that IS-PRM can be deployed on plasma samples to credential candidate plasma biomarkers for follow up validation studies. The method was capable of targeting >1,300 proteins in a highly reproducible manner, measuring >800 proteins in human plasma over several orders of magnitude with high specificity and sensitivity. Endogenous measurements across six human plasma pools included 200 proteins with reported plasma concentrations <1 ng/mL. The data demonstrate, perhaps not surprisingly^34–36^, that differences in protein expression levels between case and control pools were relatively small, highlighting the need for highly precise measurements (and perhaps multi-protein panels and longitudinal sampling of individual patients over time)^37^ to provide clinical diagnoses. IS-PRM quantification showed excellent analytical precision in both the triplicate analysis of a response curve (median CV of 7.7%) and the analysis of neighboring fractions of the human plasma pools (median CV across fractions of 11.0%), proving the method is capable of high precision.

One challenge in measuring low abundance plasma proteins is the extensive sample preparation required. In this study we incorporated abundant plasma protein depletion and bRP fractionation, which limited the analytical throughput for analyzing large numbers of samples and necessitated the use of plasma pools instead of individual plasma samples. One limitation of this study design is that a single outlier patient in one plasma pool can skew the biomarker results from that pool, a situation that can be corrected by analyzing multiple independent pools, each of which includes multiple patients, and aggregating results. This workflow was able to triage the list of candidates to a number more amenable to workflows like multiplex immuno-MRM^11,38–40^, which can be used to support clinical validation studies in a high throughput manner for the most promising candidates.

Follow-up studies will be required to determine if the PDX model serves as a viable conduit for discovering clinically translatable biomarkers. Patient-derived xenografts of human cancer have emerged as powerful tools for clinical/translational science due to their recapitulation of many aspects of the biology of tumors derived from patients, including treatment responses, genomic mutation and copy number alterations, as well as RNA and protein expression^41–43^. This high degree of biological consistency with clinical samples may make PDX-bearing mice a potential discovery platform for identification of tumor-derived proteins in plasma since human sequences can be distinguished from mouse peptides using mass spectrometry^44,45^.

Regardless of the source of biomarker candidates, IS-PRM is well suited to the challenges posed by biomarker prioritization studies, and application of the method is generally applicable to any large-scale protein quantification study. Considerations for the sample complexity, dynamic range of targeted proteins, extent of sample preparation necessary, throughput required, and the availability of internal standards may dictate which proteomic method is most appropriate for protein quantification in each application.

## Methods

### Reagents and materials

Ammonium bicarbonate (A6141), EDTA (E7889), Hydroxylamine solution (438227), Iodoacetamide (IAM; A3221), Phosphatase Cocktail 2 (P5726), Phosphatase Cocktail 3 (P0044), Protease Inhibitor (P8340), Tris (T2694), and Urea (U0631) were obtained from Sigma (St. Louis, MO). Acetonitrile (A955), TCEP (77720), and water (MS-grade; W6) were obtained from ThermoFisher Scientific (Waltham, MA). Other reagents obtained were HEPES (Alfa Aesar #J63002), EGTA (Bioworld #40520008-1), formic acid (Millipore #111670), Trypsin (Promega, V5113), Lys-C (WAKO #125-05061), and Rapigest (Waters #186002123). Heavy stable isotope-labeled standards (SIS) to 4,968 were synthesized by Vivitide (Gardner, MA) at a scale between 0.4-0.7 nmol and 25%-75% purity per peptide. The heavy peptides incorporated a fully atom labeled 13C and 15N isotope at the C-terminal K or R position of each peptide, resulting in a mass shift of +8 or +10 Da, respectively (if available; otherwise, the incorporation occurred at another amino acid residue (leucine (L), isoleucine (I), proline (P), valine (V), phenylalanine (F) or aspartic acid (D)). Peptides were solubilized in 1 mL of 3% MeCN/0.1% FA, an equal volume of each SIS peptide was mixed together and stored at −80 °C until use. Heavy standards to another 208 peptides were already available^27^ and were also included in the SIS mix.

### Identification of candidate biomarkers

All animal experiments were approved by the Baylor College of Medicine Institutional Animal Care and Use Committee (IACUC, Protocol AN-2289) and performed in compliance with the Guide for the Care and Use of Laboratory Animals of the NIH^46^. Human tumor tissue was transplanted into epithelium-free “cleared” fat pads of 4-week-old SCID/Beige (Envigo) female mice as ~1 mm^3^ fragments^47^ and allowed to grow to ~500 mm^3^. Successful transplantation of human tissue was verified by H&E staining. Blood was collected from the mouse via the inferior vena cava using a syringe filled with 50 μL 0.5M EDTA. To avoid RBC lysis, blood samples were immediately centrifuged at 2000 x g for 10 minutes. The plasma was collected into 100uL aliquots and stored at −80°C. Plasma samples were depleted of high- and mid-abundant proteins using immune-depletion columns (Agilent Multiple Affinity Removal Column MARS3 (Agilent 5188-5218) coupled to the Seppro mouse LC5 SuperMix column (Sigma S5824) or Seppro mouse LC10 IgY7 column (Sigma S5699) coupled to a Seppro mouse LC5 SuperMix column) on an ÄKTA HPLC system^48^. Protein concentrations before and after depletion were measured by BCA Protein Assay Kit (ThermoFisher, 23235). Each sample was depleted three times in succession with the depletion columns stripped, neutralized, and equilibrated between each depletion. 67 μL of mouse plasma was diluted with 433 μL of Seppro Dilution Buffer and passed through a 0.45 μm spin filter before loading onto the ÄKTA HPLC system. The depleted samples was collected in a 20 mL flowthrough fraction and the three independent depletions were pooled and concentrated in an Amicon Ultra Centrifugal Filter Unit (3 kDa cutoff; Millipore UFC900324) and buffer exchanged three times with 1x Urea buffer (6 M Urea, 25 mM Tris (pH 8.0), 1 mM EDTA, 1 mM EGTA).

Depleted plasma samples were reduced in 24 mM TCEP 30 minutes at 37 °C with shaking, followed by alkylation with 43 mM IAM in the dark at room temperature for 30 minutes. Lysates were then diluted to 2 M Urea with 200 mM Tris (pH 8.0). Lys-C was dissolved in 25 mM Tris (pH 8.0) at 200 μg/mL and added to lysates at 1:100 (enzyme:protein) ratio by mass and incubated for 2 hours at 37 °C with shaking. 200 mM Tris (pH 8.0) was added to achieve 1 M Urea final concentration and trypsin added at a 1:50 trypsin:protein ratio and incubated for 2 hours at 37°C with shaking. After 2 hours, a second trypsin aliquot was added at a 1:100 trypsin:protein ratio and incubated overnight at 37 °C with shaking. The reaction was quenched with formic acid (FA) to a final concentration 1% by volume. Samples were desalted using Oasis HLB 96-well plates (Waters 186000309) and a positive pressure manifold (Waters). The plate wells were washed with 3 × 400 μL of 50% acetonitrile (MeCN)/0.1% FA, and then equilibrated with 4 × 400 μL of 0.1% FA. The digests were applied to the wells, washed 4 x 400 μL 0.1% FA before being eluted drop by drop with 3 x 400 μL of 50% MeCN/0.1% FA. The eluates were lyophilized, followed by storage at −80 °C until use. A portion of samples were TMT-labeled according to manufacturer’s instructions prior to mass spectrometry analysis.

Digested plasma samples were fractionated using a previously described basic reverse phase chromatography workflow^49^ prior to LC-MS/MS analysis on a Thermo Scientific Orbitrap Fusion Lumos Tribrid mass spectrometer operated in positive mode. The samples were separated using a nanoACQUITY UPLC system (Waters) by reversed-phase HPLC. The analytical column was manufactured in-house using ReproSil-Pur 120 C18-AQ 1.9 um stationary phase (Dr.MaischGmbH) and slurry packed into a 25-cm length of 360 μm o.d.×75 μm i.d. fused silica picofrit capillary tubing (New Objective). The analytical column was heated to 50 °C using an AgileSLEEVE column heater (Analytical Sales and Services) and equilibrated to 98% Mobile Phase A (MP A, 3% MeCN/0.1% FA) and 2% Mobile Phase B (MP B, 90% MeCN/0.1% FA) and maintained at a constant column flow of 200 nL/min. The sample was injected into a 5-mL loop placed in-line with the analytical column which initiated the gradient profile (min:%MP B): 0:2, 1:6, 85:30, 94:60, 95:90, 100:90, 101:50, 110:50. A spray voltage of 1800 V was applied to the nanospray tip. MS/MS analysis consisted of 1 full scan MS from 350-1800 m/z at resolution 60,000 followed by data dependent MS/MS scans using 30% normalized collision energy of the 20 most abundant ions. Selected ions were dynamically excluded for 45 seconds.

### Data analysis for identification of candidate biomarkers

Raw MS/MS spectra from the analysis were searched against reviewed Mouse Universal Protein Resource (UniProt) sequence database and a combined mouse and human UniProt sequence database, release 2018_08 using MaxQuant^50^ v1.5.5.1. The search was performed with tryptic enzyme constraint set for up to two missed cleavages, oxidized methionine set as a variable modification, and carbamidomethylated cysteine set as a static modification. Peptide MH+ mass tolerances were set at 20 ppm. The overall FDR was set at ≤1%. Results from the search against the combined human/mouse databased allowed categorization of peptides into 3 classes: i. human-specific, ii. mouse-specific, and iii. ambiguous (mouse or human). All spectra that were categorized as human specific in this search that also returned an identification in the search against the mouse-only database were filtered out to ensure candidate biomarker proteins were human in origin.

### Preparation of lysates for method characterization

Yeast cells (*Saccharomyces cerevisiae*) were harvested and lysed using a previously described method^51^. MCF10A cells were obtained from American Type Culture Collection (ATCC, CRL-10317) and authenticated by short tandem repeat (STR) DNA profile. Cells were grown in DMEM:F12 (ThermoFisher 11320-033), supplemented with 5% horse serum (ThermoFisher 16050-122), 0.5 μg/mL hydrocortisone (Sigma H-0888), 20 ng/ml hEGF (ThermoFisher PHG0311), 10 μg/mL insulin (Sigma I-0516), 100 ng/mL cholera toxin (Sigma C-8052), and 100 units/mL penicillin and 100 μg/mL streptomycin (ThermoFisher 15140148) at 37 °C, 5% CO_2_. Cells were harvested at ~80% confluence by trypsinization, washed 2x with PBS and lysed in 1x Urea buffer (6 M Urea, 25 mM Tris (pH 8.0), 1 mM EDTA, 1 mM EGTA plus 1% each of Protease Inhibitor (Sigma P8340), Phosphatase Inhibitor Cocktail 2 (Sigma P5726), and Phosphatase Inhibitor Cocktail 3 (Sigma P0044) at 5×10^7^ cells / mL. Chromatin was disrupted by sonication, cleared by centrifugation (20,000 RCF, 10 minutes, 4°C) and lysates transferred to cryovials (ThermoFisher 374081) and stored in vapor phase of an LN2 tank. Digested and desalted MCF10A lysate was serially diluted with digested and desalted yeast cell lysate to make response curve concentration points that contained 100%, 10%, 1%, 0.1% and 0% MCF10A. 50 μg of each concentration point underwent SIS mix addition and bRP fractionation by the above method. 96 fractions were concatenated into 6 fractions by plate column and analyzed in triplicate

### Human plasma sample depletions, pooling, and processing

138 human plasma samples were obtained from the National Cancer Institute’s Early Detection Research Network (EDRN) biorepository^52,53^. The plasma samples were assigned to one of six pools: non-proliferative control (20 samples), proliferative control (20), atypia control (20), Her2+ (19), Triple Negative (19), and two pools of ER+Her2- (20 samples per pool). Each sample pool was further divided into four sub-pools and these sub-pools were randomized across the depletion process to reduce the chance of introducing batch effects. Plasma samples were depleted once each using human IgY14 LC10 (Sigma S5074) and human Supermix LC5 (Sigma S5324) columns coupled to an ÄKTA HPLC system^48^. Independent depleted plasma samples were collected in a single 20 mL flowthrough fraction, pooled by sub-pool, concentrated with an Amicon Ultra Centrifugal Filter Units (3 kDa cutoff, Millipore UFC900324) and buffer exchanged with 50 mM Ammonium bicarbonate (Sigma A6141). Depleted human plasma samples were denatured with 0.5% RapiGest (Waters 186002123), then digested and desalted as above. A mix of all SIS peptides was added to 200 μg of each of the six digested and desalted human plasma pools and fractionated by bRP fractionation according to the above method. 96 fractions were concatenated into 24 fractions by alternating plate column and analyzed by IS-PRM.

### High-pH reverse phase (bRP) liquid chromatography

Peptide digest was loaded onto a LC system consisting of an Agilent 1200 HPLC (Agilent, Santa Clara, CA) with mobile phases of 5 mM NH4HCO3, pH 10 (A) and 5 mM NH4HCO3 in 90% MeCN, pH 10 (B). The peptides were separated by a 2.1 mm × 250 mm Zorbax Extend-C18, 3.5 μm, column (Agilent Cat. #773700-902) over 96 minutes at a flow rate of 1.0 mL/min by the following timetable: hold 1% B for 5 minutes, gradient from 1 to 40% B for 30 minutes, 40 to 90% B for 5 minutes, hold at 90% B for 5 minutes, 90 to 1% B for 1 minute, re-equilibrate at 1% B for 14 minutes. 0.5 minute fractions were collected from 2 – 50 minutes by the shortest path by row in a 1 mL deep well plate (Thermo Cat. #95040450). Fractions were concatenated accordingly to produce 6-24 fractions for the studies described.

### Liquid chromatography tandem mass spectrometry

IS-PRM and DDA methods were implemented by LC-MS/MS on an Easy-nLC 1000 (Thermo Scientific) coupled to an Orbitrap Eclipse mass spectrometer (Thermo Scientific) operated in positive ion mode. The LC system consisted of a fused-silica nanospray needle (PicoTip™ emitter, 75 μm ID × 27 cm, New Objective) packed in-house with ReproSil-Pur C18-AQ, 3 μm and a trap (IntegraFrit™ Capillary, 100 μm ID × 2 cm, New Objective) packed with Magic C18 AQ, 5 μm, 200 Å (Bruker) with mobile phases of 0.1% FA in water (A) and 0.1% FA in 80% MeCN (B). The peptide sample was diluted in 20 μL of 0.1% FA, 3% MeCN and 4 μL was loaded onto the column and separated over 210 minutes at a flow rate of 300 nL/min with a gradient from 4 to 9% B for 2 minutes, 9 to 25% B for 78 minutes, 25 to 44% B for 60 minutes, hold 63% B for 9 minutes, 63 to 90% B for 10 minutes, hold 90% B for 1 minute. HPLC separations were carried out at 40 °C using a column heater. A spray voltage of 2200 V was applied to the nanospray tip.

### Directed DDA mass spectrometry

Directed DDA MS/MS analysis occurred over a 3 second cycle time consisting of 1 full scan MS from 300-1500 m/z at resolution 240,000 (at m/z 200), a target AGC value of 1.2e6, and maximum fill times of 50 ms followed by data dependent MS/MS scans using HCD activation with 27% normalized collision energy of the most abundant ions. The targeted mass lists consisted of 7775 entries based on m/z. Selected ions were dynamically excluded for 60 seconds after a repeat count of 1. Raw MS/MS spectra from the analysis were searched against reviewed human Universal Protein Resource (UniProt) sequence database, release 2021_01 using MSFragger^54^. The search was performed with tryptic enzyme constraint set for up to two missed cleavages, oxidized methionine and heavy labeled K, R, L, I, P, V, F and D set as a variable modification, and carbamidomethylated cysteine set as a static modification. Peptide MH+ mass tolerances were set at 20 ppm. The overall FDR was set at ≤1. Spectral library was built from the search results using SpectraST^55^. The directed DDA results and spectral library were analyzed in Skyline^56^ to identify the six most intense transitions for each precursor, and to identify the precursor charge state that had the most intense sum of these six transitions. From these results, the target precursor, six transitions and intensity thresholds (2% of the maximum height) were exported from Skyline.

### Internal standard-targeted parallel reaction monitoring (IS-PRM)

IS-PRM was adapted from the SureQuant native implementation in the instrument control software of the Orbitrap-Eclipse. The analysis occurred over a 3.5 second cycle time consisting of 6 full scan MS events from 300-1500 m/z at resolution 120,000 (at m/z 200), with each full scan corresponding to independent precursor targeted mass lists with different isolation offsets. The PRM event targeting the precursor ions selected for the heavy isotope-containing standard (“heavy”) peptides employed an Orbitrap resolution of 7,500 (at m/z 200), a target AGC value of 5e5, and maximum fill times of 10 ms. The PRM event targeting the precursor ions selected for the endogenous (“light”) peptides triggered with the detection of least 3 transitions from a group included in a transition inclusion list corresponding to the precursor ion entry in the precursor inclusion list and employed an Orbitrap resolution of 60,000 (at m/z 200), a target AGC value of 5e5, and maximum fill times of 116 ms. PRM peak integration was performed by Skyline, and the sum of all six target transitions was used for quantification. Identification was considered successful if the ratio dot product of the transition intensities between the heavy and light peptides was > 0.98. Quantification was considered successful if the PRM results contained peak areas from at least 4 transitions (light endogenous peptides) or 5 transitions (heavy SIS peptides), had at least 5 points across the peak and had a peak area greater than 5,000. All quantifications were manually checked and any one transition with interference in either the heavy or light peptide was removed from the analysis. Peptide concentrations are reported as the peak area ratio of the light and heavy peptides.

### Verification of candidate biomarkers

Peptide peak area ratio (PAR) from the individual plasma pools were filtered to include only those that were greater than two-fold greater than the maximum PAR reported in the three yeast blank samples. For the three breast cancer subtypes, PAR were compared to that in the proliferative and non-proliferative control pools. A weighted z-score for each protein was derived based on joint evidence from multiple peptides of the protein. For each peptide, a z-score was calculated by standardizing the mean intensity difference between the cancer subtype pool and the 2 control pools with the empirical standard deviation of the peptide abundances across normal pools. If there were missing results in the control pools, the standard deviation across all pools was used and the peptide z-score was weighted by 0.5. The weighted z-score for a protein was calculated as weighted sum of z-scores of peptides mapping to the protein, approximated weighted z-scores by a normal distribution and p-values were obtained from a right tailed test. The proteins were screened for markers by fitting a regression model. Relative PAR for each protein was calculated by first normalizing the peptide PAR to the median value across all plasma pools, summed together for all peptides from a given protein and regressed on the ordinal sample categories coded as: 1---non-proliferative, 2---proliferative, 3---atypia, and 4---cancer to obtain the regression coefficient and p-value of the trend for all the candidate biomarkers. Student t-tests were used to compare the mean difference between cases and controls of each of the 22 verified candidate biomarkers. The combined p-value of the 22 p-values was estimated using the meanp function of the metap R package. Permutations (n = 1000) were performed using random sets of 22 proteins.

### Public access to data

The LC-MS/MS, directed DDA, and DIA data have been deposited to the ProteomeXchange Consortium (http://proteomecentral.proteomexchange.org) via the PRIDE partner repository^25^. All PRM data have been deposited in Panorama Public^57^.

## Acknowledgements

The authors thank Kevin Schauer and Sebastien Gallien from ThermoFisher Scientific and Aaron Gajadhar from Seer for technical support in configuring the method. This research was funded by the National Cancer Institute (NCI) Early Detection Research Network (EDRN) under grant no U01CA214172, the NCI Research Specialist program (grant no. R50CA211499), and a generous donation from the Aven Foundation. The Proteomics and Metabolomics shared resource of the Fred Hutch/University of Washington Cancer Consortium is supported by NCI grant P30 CA015704. Scientific Computing Infrastructure at Fred Hutch was funded by ORIP grant S10OD028685. The content of this publication does not necessarily reflect the views or policies of the Department of Health and Human Services, nor does mention of trade names, commercial products, or organizations imply endorsement by the U.S. Government.

## Author Contributions

J.J.K., J.R.W., R.G.I., A.B., S.C., C.-F.T., T.L., C.W.L., O.M., R.L., L.A.J., P.R.G., G.L., K.D.R., S.S., J.L., P.W., and A.G.P. acquired, analyzed, or interpreted the data. G.L., K.D.R., S.S., J.L., P.W., M.T.L., A.G.P. supervised studies. J.J.K., J.R.W., A.G.P. drafted the manuscript. S.S., J.L., P.W., M.T.L., A.G.P. conceived of and designed the work.

