## Supplementary Figures for "Multiplexed triage of candidate biomarkers in plasma using internal standard triggered-parallel reaction monitoring mass spectrometry"

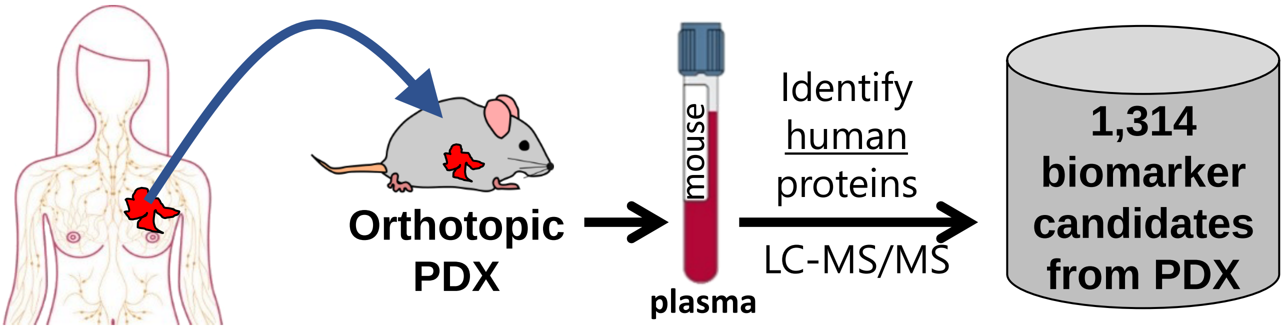


**Supplementary Fig. 1. Derivation of biomarker candidates using human tumors in mice.** Human breast cancer tumor was transplanted into SCID/Beige female mice and allowed to propagate from ~1 $mm^3$ to ~500 $mm^3$. Plasma samples from 23 PDX-bearing mice were depleted of high- and mid-abundant mouse plasma proteins by immunodepletion, pooled, proteolytically digested, fractionated by basic reverse-phase liquid chromatography, and profiled by shotgun data dependent LC-MS/MS. The resulting MS/MS data were searched against mouse and human protein sequences to identify 5,498 unique human-specific peptides in the union of the three independent profiles, mapping to 1,314 unique human proteins.

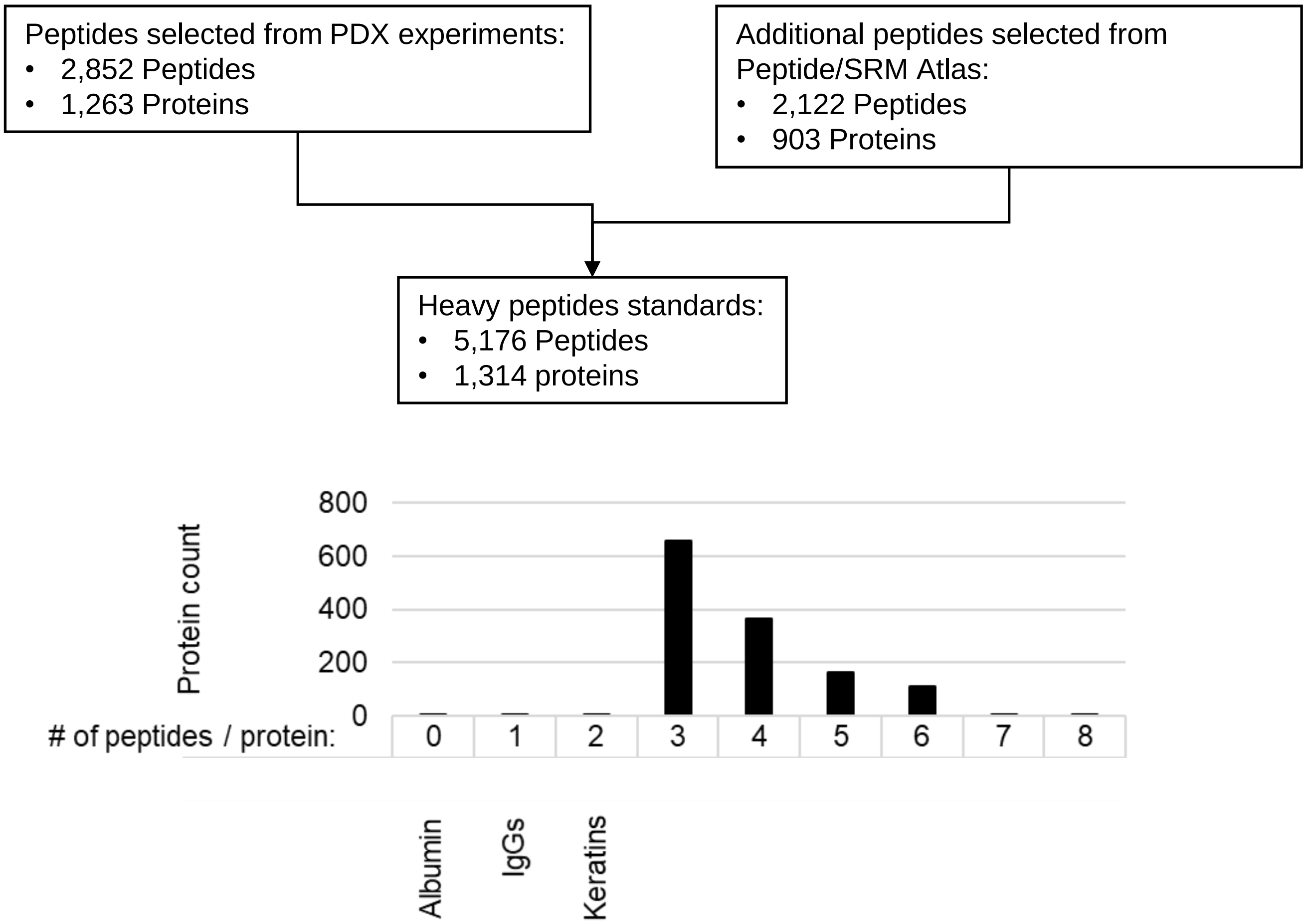


**Supplementary Fig. 2. Summary of peptides ordered for synthesis.** Peptides were selected from two sources: those directly observed in the PDX discovery experiments and the union of peptides from the online databases Peptide Atlas (http://www.peptideatlas.org/) and SRMAtlas (http://www.srmatlas.org/). For selection, peptides had to be between 7 and 25 amino acids in length; have a hydrophobicity score between 10 and 40; have no more than 1 missed cleavage. 1,303 of the candidate biomarkers are represented by three or more peptides per protein. Proteins with less than 3 peptides were keratins and IgGs. No peptides were selected for human albumin.

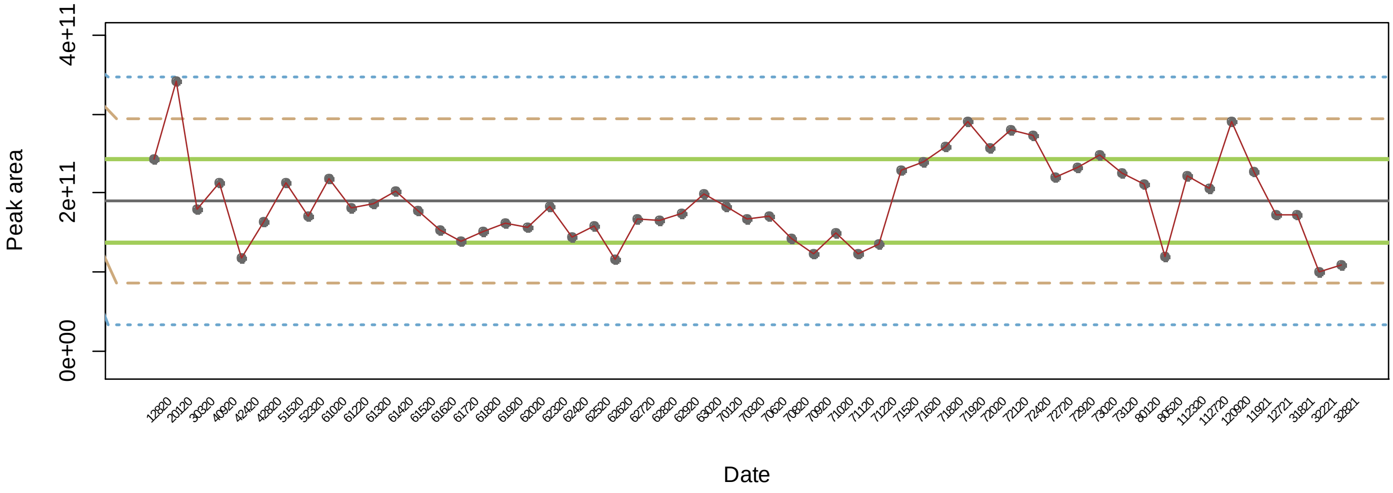


**Supplementary Fig. 4. Levey-Jennings plot of peak areas from yeast QC LC-MS/MS analysis.** The IS-PRM method relies on consistent signal intensities and mass accuracies to ensure than the internal standard triggers are applied consistently throughout the course of a study. Two quality control samples were analyzed regularly to ensure instrument stability: digested HeLa lysate analyzed every week to confirm the mass accuracy was less than 5ppm and digested yeast lysate analyzed daily to confirm signal sensitivity. For the yeast QC samples, the intensities of 11 selected peptides were compared to historical values using a Levey-Jennings plot to ensure that they passed Westgard rules. This vigorous QC program avoided any system degradation during the six months in which samples from this study were analyzed. 200 ng of a commercially available yeast lysate digest was analyzed every day that the study was run on the Thermo Eclipse instrument and the intensities of 11 selected peptides were compared to historical values using a Levey-Jennings plot to ensure that they passed Westgard rules.
